# PBX-dependent and independent Hox programs establish and maintain motor neuron terminal Identity

**DOI:** 10.64898/2025.12.24.696347

**Authors:** Manasa Prahlad, Weidong Feng, Oyunsuvd Bat-Erdene, Yihan Chen, Paschalis Kratsios

**Affiliations:** Department of Neurobiology, University of Chicago, Chicago, IL 60637, USA; Center for Motor Neuron Disease, University of Chicago, Chicago, IL 60637, USA; Neuroscience Institute, University of Chicago, Chicago, IL 60637, USA; Committee on Genetics, Genomics, and Systems Biology, Chicago, IL 60637, USA; Committee on Development, Regeneration, and Stem Cell Biology, Chicago, USA; Committee on Neurobiology, Chicago, IL 60637, USA

## Abstract

Motor neuron (MN) diversity is essential for producing the broad repertoire of animal movements, yet the molecular mechanisms that specify MN subtypes remain incompletely defined. Here, we investigate how Hox genes and their PBX cofactors shape cholinergic MN subtype identity along the anterior–posterior (A–P) axis of the *C. elegans* ventral nerve cord (VNC). In anterior MNs, we show that the anterior Hox genes *ceh-13 (Lab/Hox1)* and *lin-39 (Scr/Dfd/Hox4–5)* collaborate with the Hox cofactor *ceh-20 (Exd/Pbx1–4)* and the terminal selector *unc-3 (Collier/Ebf1–4)* to activate terminal identity genes. In posterior nerve cord MNs, the mid-body Hox gene *mab-5 (Antp/Hox6–8)* represses terminal identity gene expression by antagonizing *unc-3* in a *ceh-20*-dependent manner. Notably, *mab-5* and *ceh-20* are required not only during early development but also in later life stages to maintain posterior MN identity. In lumbar MNs, the posterior Hox gene *egl-5 (Abd-A/Abd-B/Hox9–13)* collaborates with *unc-3* to activate lumbar-specific MN terminal identity genes in a *ceh-20*-independent manner. We further find that *ceh-20* is necessary for Hox gene expression (*ceh-13, lin-39, mab-5*) in VNC MNs, supporting a model where Hox positive autoregulation requires PBX activity. Together, these findings reveal PBX-dependent and independent roles for Hox genes in establishing and maintaining MN identity, illustrating how combinatorial interactions between Hox factors and terminal selectors generate neuronal subtype diversity.

**AUTHOR SUMMARY:** Animals rely on many different types of motor neurons to generate precise and flexible movements, but how these neuron subtypes are specified remains an open question. In this study, we examine how a family of developmental genes called Hox genes, together with their cofactors, help define distinct motor neuron identities in the nervous system of the nematode *Caenorhabditis elegans*. We find that different Hox genes act in specific regions of the ventral nerve cord to either turn motor neuron identity genes on or keep them off. In anterior motor neurons, certain Hox genes work together with a cofactor called PBX and a neuron-specific regulator (UNC-3) to activate genes required for proper motor neuron function. In contrast, a mid-body Hox gene suppresses these genes in posterior neurons, while a more posterior Hox gene activates a unique set of genes in lumbar motor neurons through a different mechanism. Importantly, we show that some Hox genes and PBX are needed not only during early development but also later in life to maintain motor neuron identity. Together, our findings reveal how combinations of Hox genes and cofactors generate and preserve motor neuron diversity, providing insight into general principles of nervous system development.

## INTRODUCTION

All kinds of movement, from the fine motions we use to paint or write to the whole-body motions we use to dance and play sports, rely on the function of hundreds of muscles controlled by thousands of motor neurons (MNs). To achieve such diversity of movement, we require a corresponding diversity of MN subtypes. How this diversity is generated is an enduring question in developmental neurobiology.

In vertebrate and invertebrate animals, MN subtypes are positioned at different locations of the nervous system, displaying distinct molecular, functional, and morphological features. In quadrupeds, for example, MNs with cell bodies in the brachial region of the spinal cord innervate forelimb muscles, whereas MNs in the lumbar spinal cord innervate hindlimb muscles^1^. Similarly, MN subtypes that innervate jaw, facial, or eye muscles each occupy distinct rhombomere segments in the developing hindbrain^2^.

In addition to its well-known developmental roles in patterning the embryo along the anterior-posterior axis^3^, the highly conserved family of Hox transcription factors is known to control MN development by acting at two different levels. First, Hox genes trigger distinct downstream transcriptional programs to diversify MN progenitors in the vertebrate hindbrain^4^. In *Drosophila*, they are known to regulate neuroblast specification, proliferation, and survival ^5–7^. Second, Hox gene expression has been observed in post-mitotic MNs in *C. elegans*, *Drosophila*, zebrafish, chick, and mice, though their precise functions in these cells are still being explored ^8^ ^9^ ^10–12^ ^13–15^. Hox genes play important roles in MN survival, specification, and circuit assembly^16^. Hox depletion specifically in post-mitotic MNs is known to disrupt axon guidance and circuit assembly in *Drosophila* and mice ^17,18^. Hox proteins have also been shown to specify MN subtype identities. In *C. elegans*, *lin-39 (Scr/Dfd/Hox4-5)*, *mab-5 (Antp/Hox6-8),* and *egl-5 (Abd-B/Hox9-13)* control MN subtype terminal identity^19^. In *Drosophila*, post-mitotic RNAi depletion of Ubx and Dfd was found to disrupt locomotory and feeding MN fates, respectively ^20,21^. In mice, knockout of *Hoxa5* in post-mitotic MNs caused disorganization, irregular development, and eventually loss of phrenic MNs, resulting in respiratory failure upon birth^18^. Similarly, depletion of *Hoxc8* in post-mitotic brachial MNs resulted in dysregulation of several terminal identity genes^22^. Despite these advances, the Hox transcriptional networks required to establish and maintain MN subtype identity remain poorly understood.

Members of the Pre-B-cell leukemia transcription factor (PBX) family of homeodomain proteins are known Hox co-factors, necessary for evoking the latent DNA-binding specificity of different Hox orthologs and thus allowing them to specify distinct structures along the body axis^23^. Consequently, many Hox patterning functions depend on PBX activity^23,24^. However, some Hox functions are known to occur independently of PBX. This has been most conclusively demonstrated in *Drosophila,* where examples of PBX-independent Hox functions include haltere specification by Ultrabithorax^25^, Deformed functions in the posterior head^26,27^, and repressive functions of Ultrabithorax and Abdominal-A^28,29^. In the context of neuronal development, the PBX-dependent and independent functions of Hox proteins remain poorly defined. However, emerging evidence suggests that PBX is indeed involved in some Hox-mediated processes in MN differentiation. A study in the mouse spinal cord showed that post-mitotic depletion of *Pbx* genes in MNs affected Hox-dependent programs, disrupting the differentiation, connectivity, and organization of multiple MN subtypes^30^. Similarly, studies in the zebrafish hindbrain showed that Pbx mutant phenotypes often phenocopy Hox mutants^31–33^.

The *C. elegans* ventral nerve cord (VNC) offers an experimentally tractable system to study Hox and Pbx gene function in MN development, as the lineage, morphology, connectivity, and molecular makeup of VNC MNs are well characterized^8,34^. Molecular profiling recently showed that four of the six *C. elegans* Hox genes (*ceh-13, lin-39, mab-5, egl-5*) continue to be expressed in adult MNs of the VNC ^8^. Prior genetic studies showed that the midbody Hox genes *lin-39 (Scr/Dfd/Hox3-5*) and *mab-5 (Antp/Hox6-8)*, as well as the posterior Hox *egl-5 (Abd-A/Abd-B/Hox9-13)* control distinct terminal identity features of midbody and posterior MNs, respectively ^9,19,35^. However, the role of the anterior Hox gene *ceh-13* (*Lab/Hox1*) in MN development remains unclear due to the early lethality of *ceh-13* mutants^36^. Further, adult-specific depletion of LIN-39 causes loss of expression of acetylcholine biosynthesis genes, suggesting that LIN-39 is required to maintain adult MN terminal identity^35^. Whether other HOX and PBX factors are similarly required in adult stages remains untested.

Recent studies in *C. elegans* revealed that Hox factors collaborate with terminal selector-type TFs, which determine the identity and function of individual neuron types throughout life^37–38^. UNC-3, the sole *C. elegans* ortholog of the Collier/Olf/Ebf (COE) family of TFs, functions as a terminal selector of VNC cholinergic MNs by directly activating a large battery of terminal identity genes (e.g., acetylcholine biosynthesis components, ion channels, neuropeptides) ^39, 40, 41^. The midbody Hox protein LIN-39 collaborates with UNC-3 in cholinergic MNs located in the midbody region of the VNC, whereas the posterior Hox protein EGL-5 collaborates with UNC-3 in lumbar cholinergic MNs^19,35^. Similarly, UNC-30/PITX, the terminal selector of GABAergic MN identity^42,43^, works together with LIN-39 in GABAergic MNs located in the midbody region of the VNC^44^. Hence, the intersectional activity of region-specific TFs and neuron type-specific TFs (terminal selectors) is needed to determine the identity of distinct MN subtypes located at different positions along the anterior-posterior (A–P) axis of the *C. elegans* VNC (**Fig. 1A**). However, the underlying molecular mechanisms, including the involvement of PBX factors, remain largely unknown.

**Figure 1:**
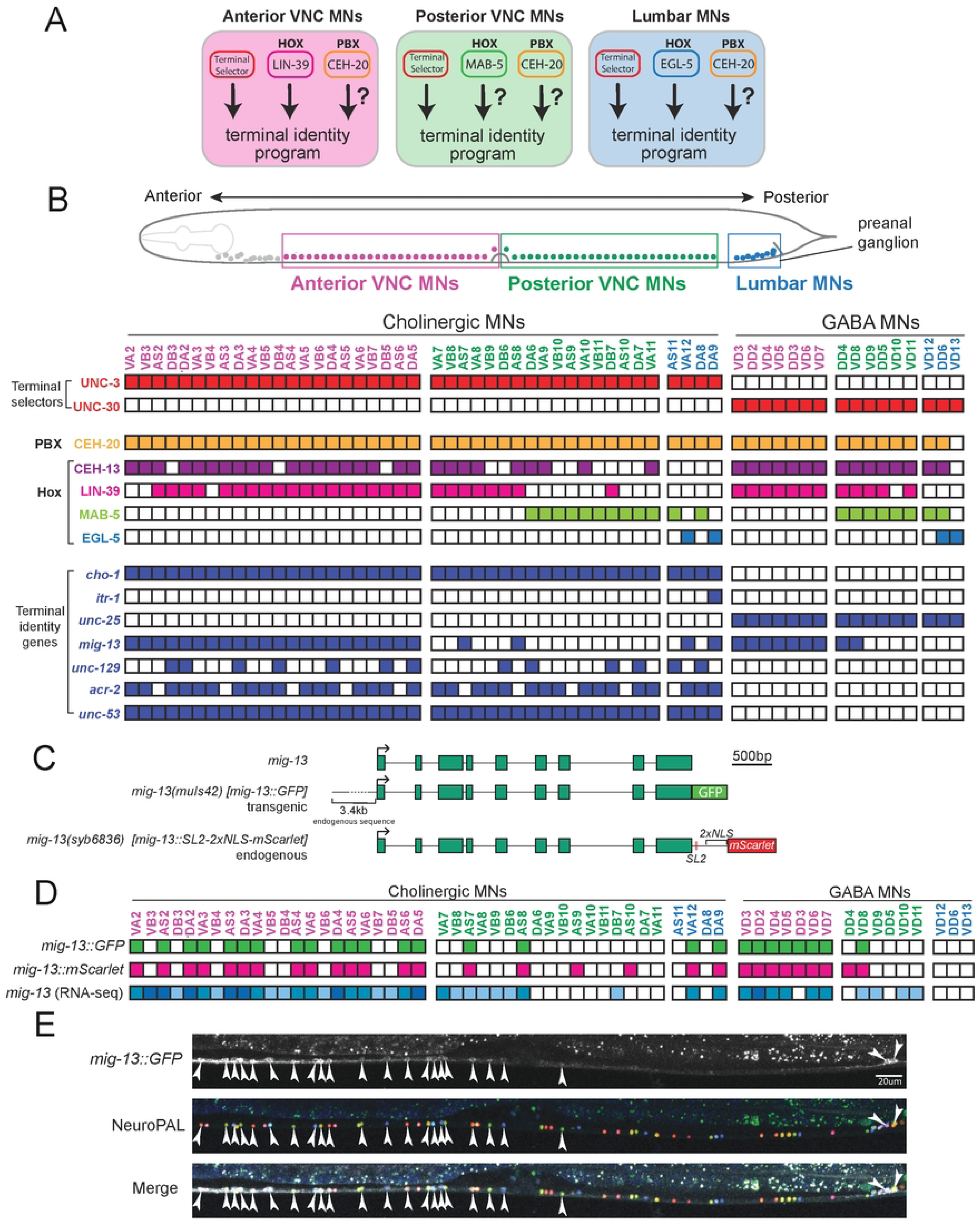
The *C. elegans* ventral nerve cord as a model to study neuronal diversification. **A:** Model depicting co-expression of *unc-3*, Hox genes and *ceh-20* in different MN subtypes. Each panel represents MNs in different regions of the A-P axis. Question marks indicate that the function of Hox and *ceh-20/PBX* in MN terminal identity remains unclear. **B:** Top: schematic showing MNs occupying three different regions along the A-P axis: anterior VNC (magenta), posterior VNC (green), and preanal ganglion (blue). Bottom: TF and key terminal identity gene expression patterns shown with single-cell resolution. **C:** Schematics of endogenous *mig-13* locus, *muIs42[mig-13::GFP]* transgenic translational reporter containing 3.4kb endogenous upstream sequence, and endogenous *mig-13* (*syb6836[mig-13::SL2::2xNLS::mScarlet])* transcriptional reporter. **D:** Single-cell expression patterns of *muIs42* and *syb6836* in L4 MNs, as well as *mig-13* transcript levels in regional subclasses identified from scRNA-seq data in adult (day 1) MNs^8^. **E:** Representative images of (L4 stage) of *muIs42[mig-13::GFP]* animals with NeuroPAL.

In this study, we unveil PBX-dependent and independent functions of Hox genes that determine MN terminal identity in *C. elegans*. In cholinergic MNs of the anterior VNC, we find that *ceh-13 (Lab/Hox1), lin-39 (Scr/Dfd/Hox3-5*) and *ceh-20 (Exd/Pbx1-4)* collaborate with the terminal selector *unc-3* to promote anterior MN terminal identity. In MNs of the posterior VNC, the midbody Hox gene *mab-5 (Antp/Hox6-8)* represses the expression of anterior MN identity genes by antagonizing *unc-3* in a *ceh-20 (Exd/PBX)*-dependent manner. Intriguingly, we find that MAB-5 and CEH-20 are not only required during development, but also in late larval stages to maintain posterior MN identity. Further, we find that *ceh-20* is required for Hox (*ceh-13, lin-39, mab-5*) gene expression in VNC MNs and propose a model where Hox positive autoregulation in these cells requires PBX activity. In lumbar cholinergic MNs, the posterior Hox gene *egl-5 (Abd-A/Abd-B/Hox9-13)* and *unc-3* collaborate to activate lumbar-specific MN terminal identity genes in a *ceh-20 (Exd/PBX)*-independent manner. Altogether, our findings showcase PBX dependent and independent roles of Hox genes in establishing and maintaining MN subtypes along the A-P axis of the *C. elegans* VNC, enhancing our mechanistic understanding of neuronal diversification processes.

## RESULTS

### The *C. elegans* ventral nerve cord as a model to study motor neuron diversification

In the *C. elegans* VNC, a structure analogous to the vertebrate spinal cord, MNs are divided into two groups based on neurotransmitter usage: cholinergic and GABAergic MNs (**Fig. 1B**). Based on anatomical criteria, cholinergic MNs are further subdivided into six classes (DA, DB, VA, VB, AS, VC) and GABAergic MNs into two (DD, VD) (**Fig. 1B**). Individual MNs from each class intermingle along the VNC and its flanking ganglia. All cholinergic nerve cord MNs express sets of “shared” genes that encode enzymes and receptors necessary for acetylcholine biosynthesis (e.g.,*cho-1/ChT, unc-17/VAChT*). Similarly, all GABAergic MNs express *unc-25/GAD* and *unc-47/VGAT*, both involved in GABA biosynthesis (**Fig. 1B**). However, prior work has shown that in addition to the shared genes common to all classes of cholinergic or GABAergic MNs, there are genes whose expression is restricted to MNs located in specific positions along the A-P axis^8,19^. Herein, we refer to these as “subclass-specific” genes. For example, *itr-1*/inositol triphosphate receptor, a gene encoding a calcium ion channel, is specifically expressed in DA9, a cholinergic MN in the preanal ganglion (**Fig. 1B**).

A notable subclass-specific gene is *mig-13*, ortholog of human Low-density lipoprotein Receptor-related Protein 12 (LRP12), previously reported to be expressed in both cholinergic and GABAergic MNs^45^. We sought to establish the expression pattern of *mig-13* with single-cell resolution and found it to be region-specific in the in the VNC. Using animals carrying either a transgenic *mig-13::GFP* reporter (**Fig. 1C**), or an endogenous *mig-13::SL2::NLS::mScarlet* reporter (**Fig. 1C**, see Methods), we observed a near complete overlap between the two reporters at the L4 stage, as well as with available single-cell RNA-sequencing data from day 1 adult MNs (**Fig. 1D**). Specifically, in VNC MNs anterior to the vulva, both *mig-13* reporters are expressed in most cholinergic MNs and all GABAergic cells (**Fig. 1D and E**). Posterior to the vulva, both reporters are expressed sparsely (2-4 cholinergic and 1-2 GABAergic MNs) (**Fig. 1D and E, Suppl. Fig 1**). In MNs of the preanal ganglion, referred herein as lumbar MNs, each *mig-13* reporter is expressed in two cholinergic MNs (DA9, VA12) (**Fig. 1D and E**). This region-specific expression pattern of *mig-13* in MNs offers an entry point to study the gene regulatory mechanisms that generate MN diversity along the A-P axis of the *C. elegans* ventral nerve cord.

### The terminal selectors UNC-3 and UNC-30 respectively control *mig-13* in cholinergic and GABAergic MNs

Since *mig-13* is expressed both in cholinergic and GABAergic MNs, we reasoned that it is controlled by the terminal selectors UNC-3/EBF and UNC-30/PITX, respectively (**Fig. 1B**). Using an *unc-3(n3435)* loss-of-function allele^46^, we observed a partial loss of *mig-13::GFP* specifically in MNs of the anterior VNC (**Fig. 2A**). Loss of *unc-3* had no effect on *mig-13::GFP* expression in posterior VNC MNs. Using NeuroPAL, a polychromatic fluorescent reporter strain that allows for nervous system-wide cell identification^47^, we found that only a subset of cholinergic MNs in the anterior VNC lost *mig-13::gfp* expression (**Fig. 2A**). Following a similar strategy, we observed a partial loss of *mig-13::GFP* expression specifically in GABAergic MNs of the anterior VNC in homozygous animals carrying the *unc-30(e191)* loss-of-function allele^48^ (**Fig. 2B**). In lumbar MNs, we observed loss of *mig-13::GFP* expression in DA9 and VA12 MNs in homozygous *unc-3(n3435)* animals (**Fig. 2C**), consistent with prior work ^19^. Altogether, this analysis revealed partial and region-specific effects for the terminal selectors *unc-3* and *unc-30*, suggesting that additional factors with spatially restricted activity (X, Y, Z in **Fig. 2**) are involved in the control of *mig-13*. For the ensuing analysis, we solely focus on cholinergic MNs.

**Figure 2:**
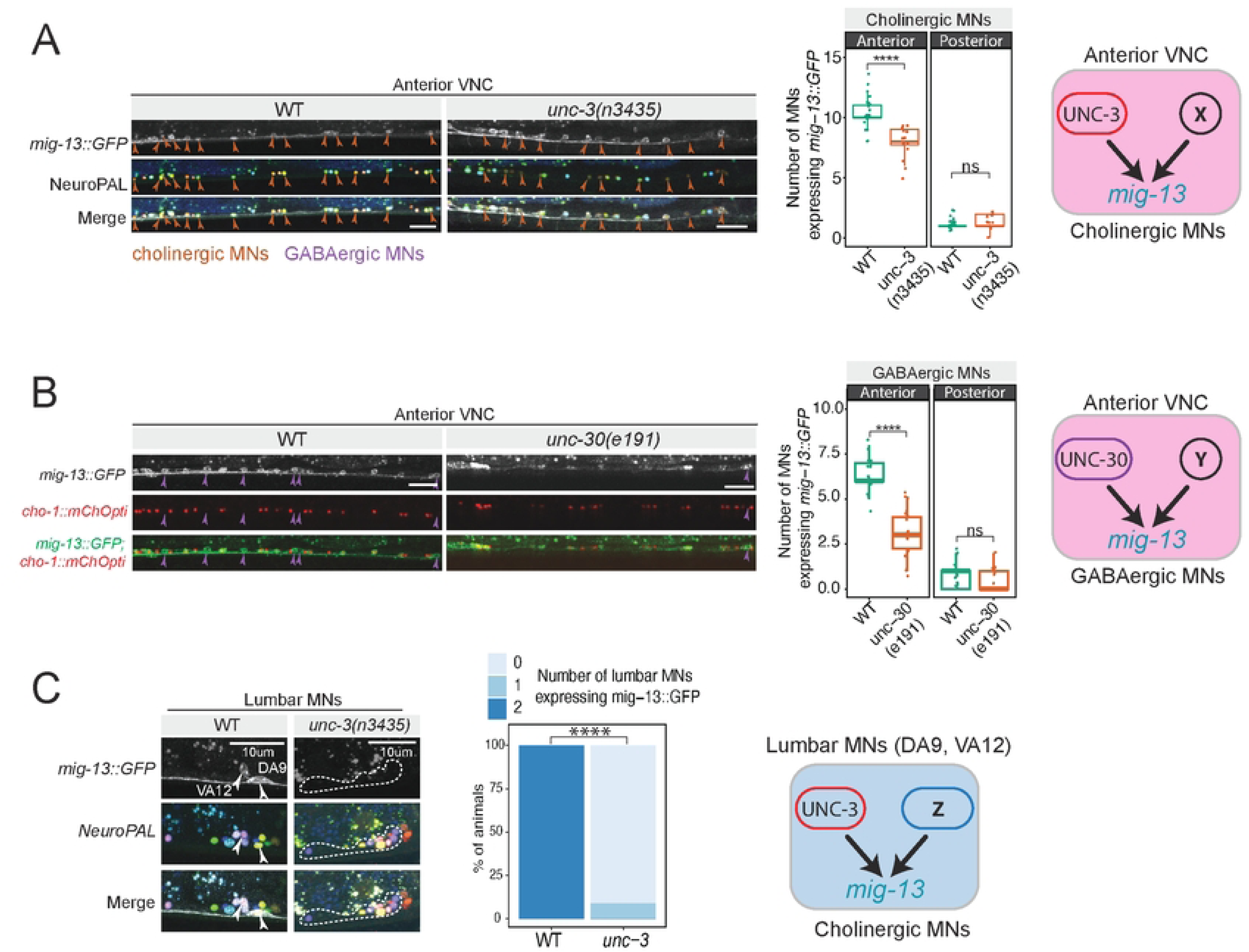
The terminal selectors UNC-3 and UNC-30 respectively control *mig-13* in cholinergic and GABAergic MNs. **A:** Left: In *unc-3* mutant animals, only some cholinergic cells lose *mig-13* expression, suggesting that UNC-3 works alongside a co-activator to drive *mig-13* expression in these cells. Middle: quantification of data. Right: schematic of *mig-13*’s regulation in cholinergic cells. Quantification of MN number was performed at the L4 stage. n > 16. For all quantifications, unpaired two-sided Welch’s t-test was performed, p<.05 = *; p<.01 = **; p<.001 = ***; p<.0001 = ****. Box and whisker plots were used; all data points presented. Box boundaries indicate the 25th and 75th percentile. The limits indicate minima and maxima values. Centre values (mean) are highlighted with a horizontal line. **B:** In *unc-30(-)* animals, only some GABAergic MNs lose *mig-13*::GFP expression, suggesting UNC-30 also works with a co-activator to activate *mig-13*. Quantified in middle panel (N= 18), schematized in right. **C:** In the lumbar motor neurons of the posterior ganglion, DA9 and VA12 express *mig-13* in WT animals. In *unc-3* mutants, both MNs consistently lose expression of *mig-13*, demonstrating that *unc-3* activity is required for *mig-13* activation in these cells. Quantification shown in middle panel, N > 17. Schematic shown on right.

### The anterior Hox gene *ceh-13* (*Lab/Hox1*) controls anterior MN identity

Four of the six *C. elegans* Hox genes (*ceh-13, lin-39, mab-5, egl-5*) are expressed continuously, from development through adulthood, in MNs^8^. Their expression patterns have been established with single-cell resolution ^8,19^, offering a tractable system to understand how Hox genes and the terminal selector UNC-3 control diverse MN identities along the A-P axis of the VNC (**Fig. 1B**).

The role of the anterior Hox gene *ceh-13* (*Lab/Hox1*) in MNs has remained unknown, partly due to the early larval lethality of *ceh-13* mutants^36^. The expression of *ceh-13* significantly overlaps with *mig-13* in MNs of the anterior VNC (**Fig. 1B**)^8^. Therefore, we evaluated *mig-13* expression in *ceh-13 (sw1)* loss-of-function mutants. At the first larval stage (L1), we observed a significant loss of *mig-13::GFP* in MNs of the anterior VNC (**Fig. 3A**). Similarly, *ceh-13* loss led to reduced expression of three additional terminal identity markers (*unc-53/NAV1, unc-129/TGFbeta,* and *acr-2/AChR)* (**Fig. 3B**), uncovering a critical requirement for *ceh-13* in anterior MN terminal identity. Importantly, we observed the same number of neurons (labeled with a panneuronal *rab-3::tagRFP* reporter) in the VNC of control and *ceh-13 (sw1)* animals (**Fig. 3C**), indicating that loss of *ceh-13* does not impact MN generation.

**Figure 3:**
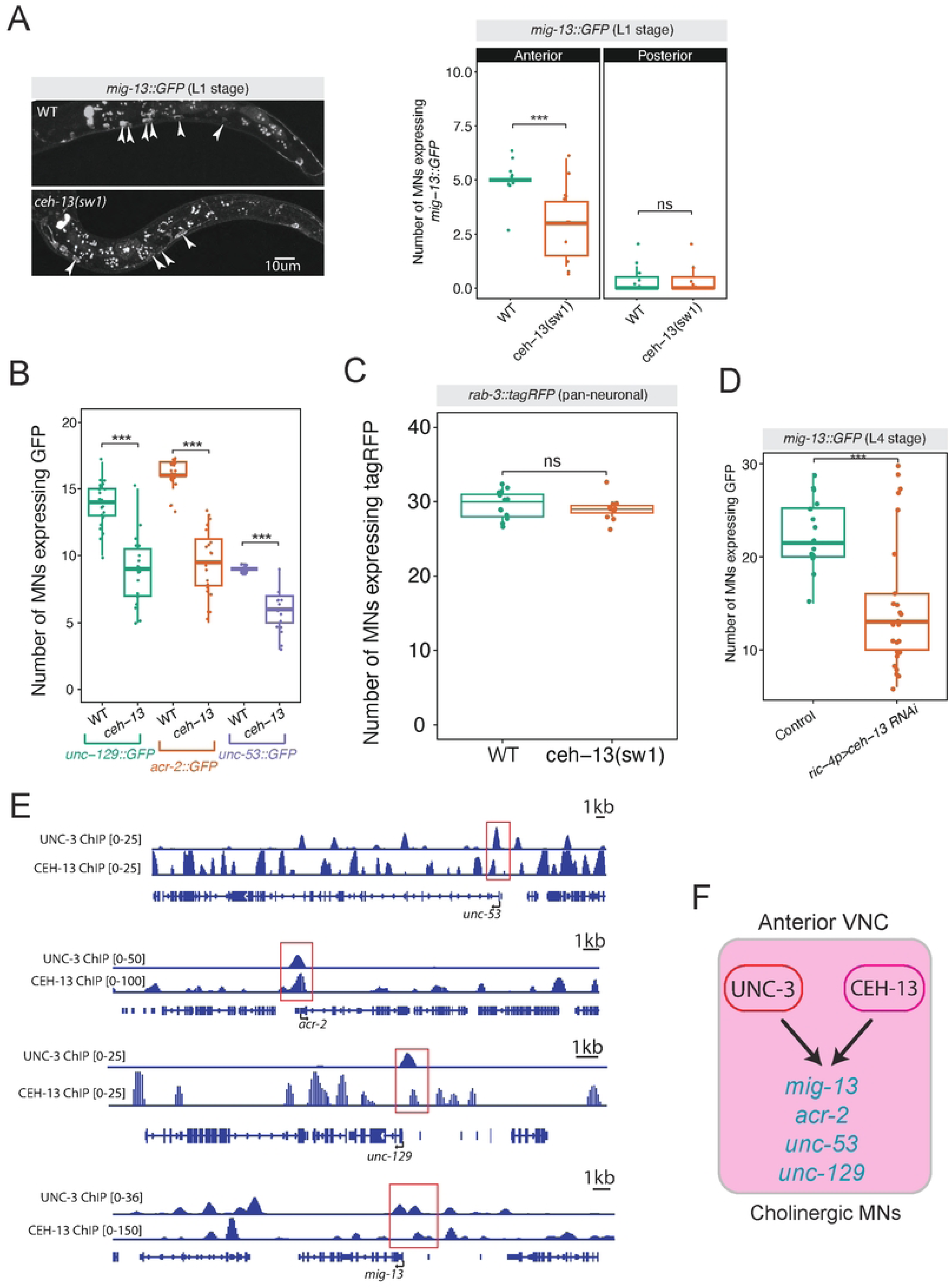
The anterior Hox gene *ceh-13* (*Lab/Hox1*) controls anterior MN identity. **A:** *ceh-13(sw1)* animals show partial loss of *mig-13* in anterior VNC MNs. Quantification shown in right panel. Quantification of MN number was performed at the L1 stage. n > 10. For all quantifications, unpaired two-sided Welch’s t-test was performed, p<.05 = *; p<.01 = **; p<.001 = ***; p<.0001 = ****. Box and whisker plots were used; all data points presented. Box boundaries indicate the 25th and 75th percentile. The limits indicate minima and maxima values. Centre values (mean) are highlighted with a horizontal line. **B:** Compared to control animals, *ceh-13(sw1)* mutants show decreased expression of *unc-129, acr-2,* and *unc-53* reporters in nerve cord MNs at L1. N > 16. **C:** *rab-3::tagRFP* expression is unaffected in *ceh-13(sw1)* animals at L1, suggesting that *ceh-13* loss does not affect pan-neuronal gene expression, nor MN generation. **D:** MN-specific RNAi against *ceh-13* causes loss of *mig-13* expression in nerve cord MNs at the L4 stage. N > 20 **E:** CEH-13 ChIP-seq shows binding peaks in each of these terminal identity gene loci, in proximity to UNC-3 ChIP-Seq peaks. **F:** Model showing CEH-13 and UNC-3 collaborating to activate terminal identity genes in cholinergic MNs.

Next, we wondered whether *ceh-13* controls MN identity in a cell-autonomous manner. To test this, we conducted MN-specific RNAi against *ceh-13* using a previously characterized promoter fragment of *ric-4* ^49^ and observed reduced *mig-13::GFP* expression in MNs (**Fig. 3D**), suggesting a cell-autonomous mode of action. Analysis of available ChIP-Seq data^50^ showed CEH-13 binding in the *cis*-regulatory regions of *mig-13, unc-53, unc-129*, and *acr-2* (**Fig. 3E**), suggesting that CEH-13 acts directly to activate their expression. Because UNC-3 is also known to directly activate these four genes ^39,51^, we propose that CEH-13 and UNC-3 collaborate to directly activate terminal identity programs in cholinergic MNs of the anterior VNC (**Fig. 3E-F**).

### The mid-body Hox gene *lin-39* (*Scr/Dfd/Hox3-5*) controls *mig-13* in anterior MNs

Like *ceh-13* (*Lab/Hox1*), the mid-body Hox gene *lin-39* (*Scr/Dfd/Hox3-5*) is expressed in anterior VNC MNs ^44^, where it co-regulates with UNC-3 several terminal identity genes, including *unc-53/NAV1, unc-129/TGFbeta,* and *acr-2/AChR*^19,35,44^. We found that *mig-13::*GFP expression in anterior MNs is partially affected in animals carrying a *lin-39(n1760)* loss-of-function allele^52^(**Fig. 4A-B**), extending the repertoire of LIN-39 target genes (**Fig. 4C**). Using the NeuroPAL strain, we found that loss of *mig-13::*GFP expression is specific to cholinergic MNs (DA, DB, VA, VB, and AS classes), as GABAergic MNs were unaffected (**Fig. 4B**). The loss of *mig-13::GFP* expression in anterior VNC MNs of *lin-39* mutants is similar to the effect observed in *ceh-13* and *unc-3* mutants (**Fig. 2C, 3A**), suggesting that two different Hox genes, *lin-39* and *ceh-13,* collaborate with the terminal selector *unc-3* to activate the terminal identity program in cholinergic MNs of the anterior VNC (**Fig. 4C**). Supporting a direct mode of gene activation, available ChIP-Seq data show that LIN-39, like CEH-13 and UNC-3, binds to the *cis*-regulatory region of *mig-13* (**Suppl. Fig. 2**).

**Figure 4:**
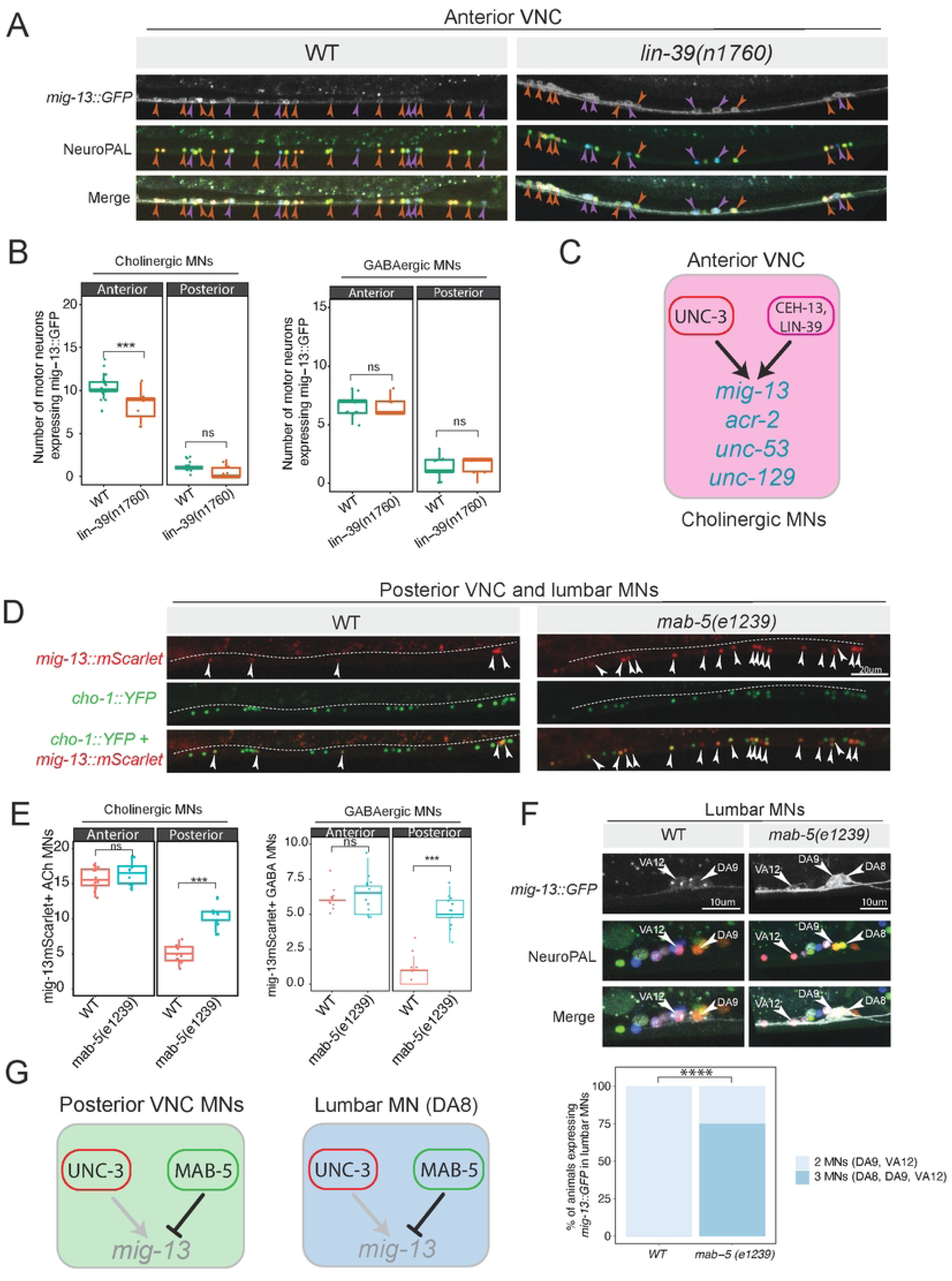
The mid-body Hox gene *lin-39* (*Scr/Dfd/Hox3-5*) and *mab-5 (Antp/Hox6-8)* control *mig-13* in anterior MNs. **A:** Compared to WT animals at L4, *lin-39* mutants have fewer MNs expressing *mig-13::GFP*. This partial effect in anterior VNC MNs suggests that *lin-39* activity is redundant with that of *unc-3* and/or *ceh-13*. **B:** Quantification of MNs expressing *mig-13::GFP* in WT and *lin-39(n1760)* mutants. Quantification of MN number was performed at the L4 stage. n > 15. For all quantifications, unpaired two-sided Welch’s t-test was performed, p<.05 = *; p<.01 = **; p<.001 = ***; p<.0001 = ****. Box and whisker plots were used; all data points presented. Box boundaries indicate the 25th and 75th percentile. The limits indicate minima and maxima values. Centre values (mean) are highlighted with a horizontal line. Left: in cholinergic MNs, right: GABAergic MNs. **C:** Schematic of *mig-13* regulation in cholinergic MNs of the anterior VNC. **D:** In *mab-5(e1239)* animals, *mig-13::mScarlet* is ectopically expressed in both cholinergic and GABAergic MNs of the posterior VNC (L4). **E:** Quantification of data shown in **D**. Quantification of MN number was performed at the L4 stage. n > 15. For all quantifications, unpaired two-sided Welch’s t-test was performed, p<.05 = *; p<.01 = **; p<.001 = ***; p<.0001 = ****. Box and whisker plots were used; all data points presented. Box boundaries indicate the 25th and 75th percentile. The limits indicate minima and maxima values. Centre values (mean) are highlighted with a horizontal line. Left: in cholinergic MNs, right: GABAergic MNs. **F:** *mab-5(e1239)* animals show ectopic *mig-13::GFP* expression in the lumbar MN DA8. DA8 was identified with the NeuroPAL strain (middle panel) and its distinct axonal trajectory. Quantification is provided below. N > 15, Fisher’s exact test: **** p = 2.2e-16. **G:** Schematics showing *mig-13* regulation in posterior VNC and lumbar (DA8) MNs.

### *mab-5 (Antp/Hox6-8)* represses *mig-13* expression in posterior VNC motor neurons

The midbody Hox gene *mab-5 (Antp/Hox6-8)* is selectively expressed in posterior VNC MNs^8^, which mostly lack *mig-13* (**Fig. 1B**). Intriguingly, we observed ectopic expression of *mig-13::GFP or mig-13::mScarlet* in MNs of the posterior VNC in animals carrying a strong loss-of-function allele for *mab-5(e1239)*^53^, indicating that *mab-5* represses *mig-13* in these cells (**Fig. 4D, Suppl. Fig 3**). The mode of repression is likely direct; ChIP-Seq shows that MAB-5 binds directly to the *mig-13* locus (**Suppl. Fig. 2**). Next, we asked whether this ectopic expression occurred in cholinergic or GABAergic MNs. To address this, we used a cholinergic MN-specific marker (*cho-1::YFP)* and evaluated expression of the endogenous *mig-13::mScarlet* reporter in both WT and *mab-5(e1239)* animals. We found that *mig-13* is ectopically expressed in both cholinergic and GABAergic MNs of the posterior VNC (**Fig. 4E**). Importantly, loss of *mab-5* does not affect the cholinergic identity of posterior VNC MNs, as the expression of *cho-1::YFP* remained unaltered (**Fig. 4E**).

Posterior to the VNC, the cholinergic lumbar MN DA8 of the posterior ganglion also expresses *mab-5*^8^ and does not express *mig-13::GFP* in WT animals (**Fig. 1D**). In *mab-5(e1239)* mutants, we found that *mig-13::GFP* is ectopically expressed in DA8 (**Fig. 4F**), consistent with prior work suggesting that MAB-5 is required to establish a unique DA8 identity distinct from other DA MNs^9,19^.

Altogether, these findings support the idea that Hox genes exert distinct, region-specific effects on MN terminal identity along the A-P axis. In MNs of the anterior VNC, *lin-39* and *ceh-13* collaborate with the terminal selector *unc-3* to activate terminal identity genes (**Fig. 4C**). In MNs of the posterior VNC and DA8, *mab-5* represses terminal identity gene *(mig-13*) expression despite the presence of *unc-3* (**Fig. 4G**).

### MAB-5 antagonizes UNC-3 to repress *mig-13* expression in posterior VNC MNs

UNC-3 is known to act as a transcriptional activator of MN terminal identity genes^39^, and is required for *mig-13* activation in MNs of the anterior VNC (**Fig. 2A**). To determine whether the ectopic *mig-13::GFP* expression in posterior VNC MNs of *mab-5* mutants requires *unc-3*, we generated *unc-3(-); mab-5(-)* double mutants that also carry NeuroPAL. We indeed observed a reduction in the number of MNs ectopically expressing *mig-13*::GFP compared to *mab-5* single mutants. NeuroPAL analysis confirmed that this reduction occurred only in cholinergic MNs of *unc-3(-); mab-5(-)* mutants (**Fig. 5A-B**). However, the reduction is partial, i.e., some cholinergic MNs in the posterior VNC of *unc-3(-); mab-5(-)* mutants continue to ectopically express *mig-13::GFP* compared to WT animals (**Fig. 5A-B**), suggesting that when *mab-5* activity is disrupted, *unc-3* is able to activate *mig-13* in posterior VNC MNs, but additional, yet-unknown factors must also be involved to drive ectopic *mig-13* expression. To test this possibility, we considered *lin-39,* which, like *unc-3*, is known to act as a transcriptional activator of terminal identity genes in VNC MNs ^19,35,44^. We found a modest reduction in the number of posterior MNs ectopically expressing *mig-13* expression in *lin-39(-); mab-5(-)* double mutants compared to *mab-5(-)* single mutants (**Suppl. Fig. 3**). Altogether, we conclude that *unc-3* and *lin-39* can activate *mig-13* in posterior MNs of the VNC when *mab-5* activity is disrupted (**Fig. 5C**).

**Figure 5:**
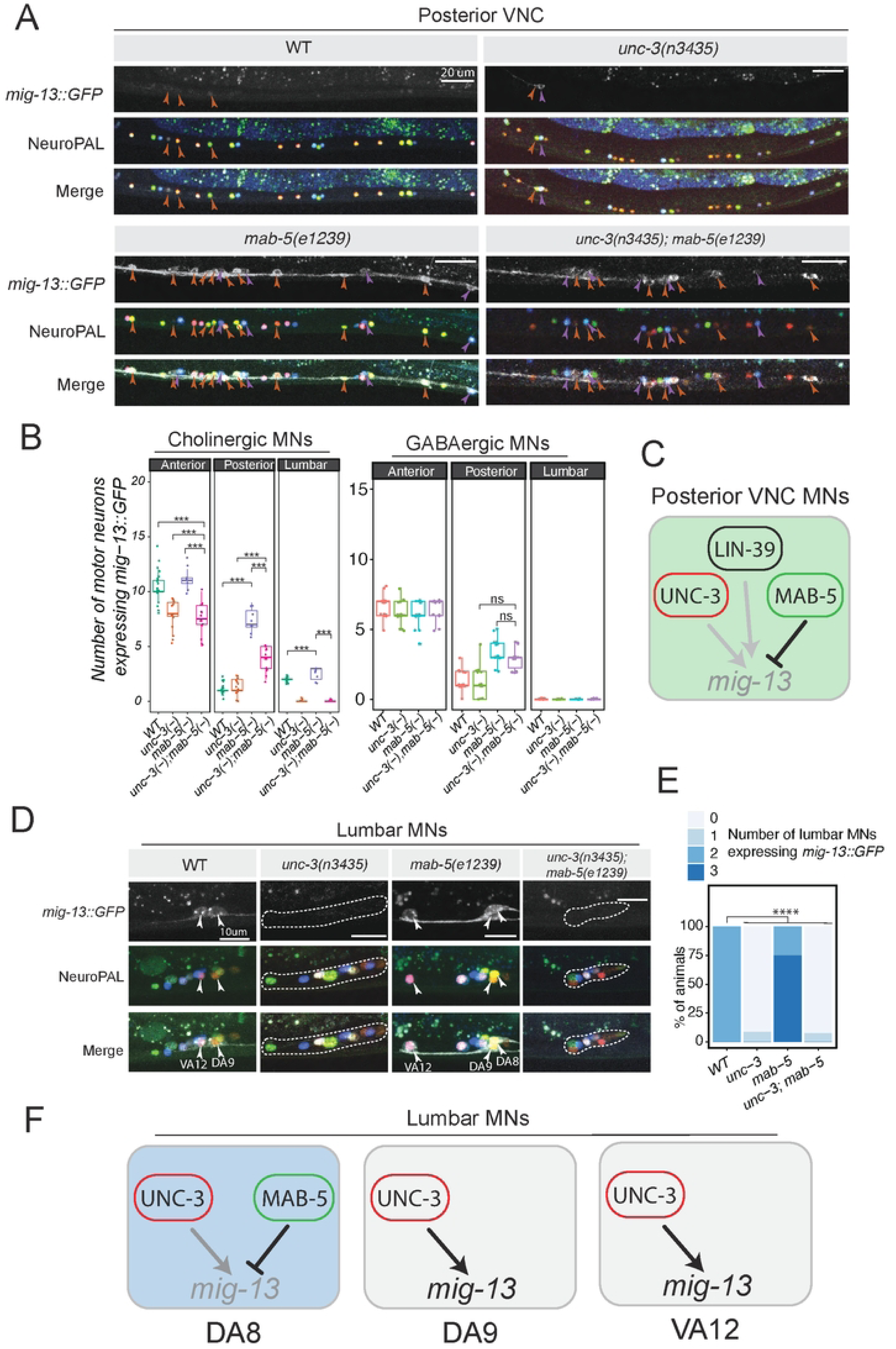
MAB-5 antagonizes UNC-3 to repress *mig-13* expression in posterior MNs. **A:** *mig-13*::GFP expression in MNs of *unc-3(-)*, *mab-5(-)*, and *unc-3(-);mab-5(-)* animals. *unc-3* activity is required for ectopic *mig-13::gfp* expression in posterior cholinergic MNs. Lack of *unc-3* does not affect *mig-13* expression in GABAergic MNs. **B:** Quantification of *mig-13*::GFP expression in cholinergic (left) and GABAergic MNs (right). Quantification of MN number was performed at the L4 stage. n = 16. For all quantifications, unpaired two-sided Welch’s t-test was performed, p<.05 = *; p<.01 = **; p<.001 = ***; p<.0001 = ****. Box and whisker plots were used; all data points presented. Box boundaries indicate the 25th and 75th percentile. The limits indicate minima and maxima values. Centre values (mean) are highlighted with a horizontal line. **C:** Schematic of *mig-13* regulation in posterior VNC MNs. MAB-5 represses *mig-13* by antagonizing UNC-3. **D-E:** The ectopic expression of *mig-13::GFP* in DA8 (lumbar MN) of *mab-5(e1239)* mutants also depends on *unc-3*. Lumbar MNs (DA8, DA9, VA12) were identified using the NeuroPAL strain. Quantification (E). N > 15. **F:** Models of *mig-13* regulation in lumbar MNs (DA8, DA9, VA12).

In the lumbar MN DA8, we witnessed similar effects: the ectopic *mig-13::GFP* expression in DA8 in *mab-5(-)* animals is lost in *unc-3(-); mab-5(-)* double mutant animals (**Fig. 5D-E**). Collectively, these findings suggest that MAB-5 antagonizes the ability of UNC-3 to activate *mig-13* in posterior VNC and lumbar DA8 MNs. Consistently, *mab-5* is not expressed in two other lumbar MNs, DA9 and VA12, in which *unc-3* is able to activate *mig-13* (**Fig. 5D-F**).

### CEH-20/PBX-dependent and independent functions of Hox genes in MNs

To gain mechanistic insights, we wondered about the involvement of PBX proteins, which are known Hox cofactors during early animal development^54,55^. The *C. elegans* genome contains three orthologs of the PBX family: *ceh-20, ceh-40,* and *ceh-60*. In this study, we focus on *ceh-20 (Exd/Pbx1–4)*, as it is expressed robustly in *C. elegans* MNs compared to *ceh-40* and *ceh-60*, and is the only one whose expression in MNs persists into adulthood^8,56^. To determine whether *ceh-20* is involved in MN terminal identity, we sought to establish its expression pattern with single-cell resolution. To this end, we generated with CRISPR/Cas9 an endogenous *ceh-20(syb6724)* reporter allele that carries the *mNG::3xFLAG::AID* cassette at the C-terminus (**Fig. 6A**). Using a cholinergic MN marker (*cho-1::mChOpti*) in larval stage 4 (L4) animals, we found *ceh-20::mNG::3xFLAG::AID* expression in all MNs of the VNC and most lumbar MNs (**Fig. 6B-C, Suppl. Fig 4**).

**Figure 6:**
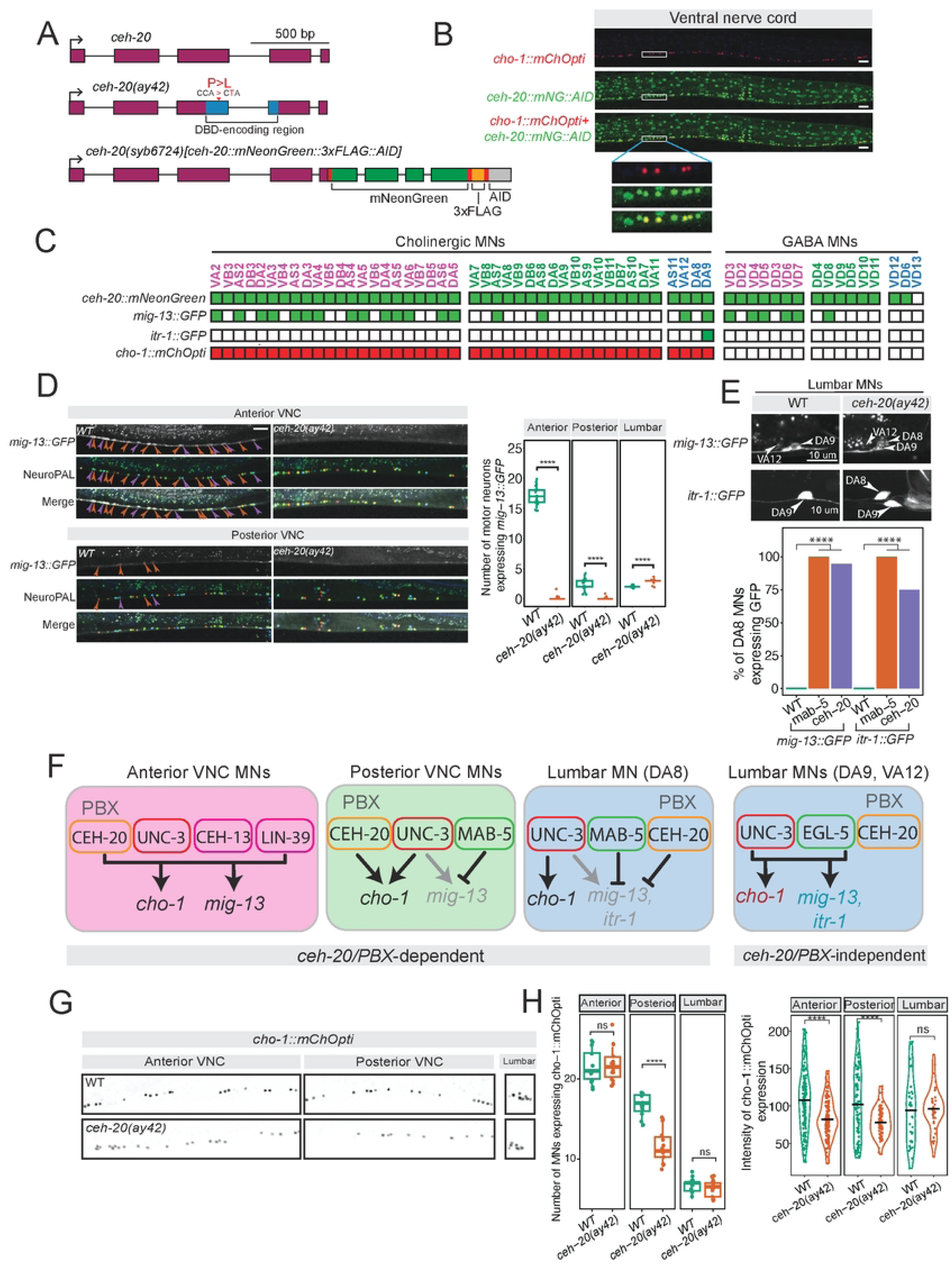
CEH-20/PBX regulates terminal identity genes. **A:** *ceh-20* locus with alleles used: *ceh-20(ay42)* hypomorph, with a point mutation in DBD, and *ceh-20*(*syb6724*) with mNeonGreen, 3xFLAG, and AID tags inserted before the stop codon. Linker sequences highlighted in red. **B:** Expression of endogenous *ceh-20(syb6724)[ceh-20::mNG::3xFLAG::AID]* reporter with cholinergic MN marker *cho-1::mChOpti*. ceh-20 is expressed in all VNC MNs, both cholinergic and GABAergic, at the L4 stage. **C:** Single-cell expression of *ceh-20::mNG::AID* in VNC and lumbar MNs. Expression of *mig-13, itr-1,* and *cho-1* reporters is shown. **D**: Loss of *mig-13*::GFP expression in nerve cord MNs (anterior and posterior) of *ceh-20(ay42)* animals. Quantification on the right. **E**: The terminal identity markers *mig-13* and *itr-1* (DA9 marker) are ectopically expressed in DA8 of *ceh-20(ay42)* mutants. Quantification shown in bottom panel. N > 15, Fisher’s exact test: p < 2.2e-16 for both genotypes. **F**: Schematic summary of our findings in VNC and lumbar MNs. **G-H:** Decreased levels of cholinergic marker *cho-1* expression in VNC MNs of *ceh-20(ay42)* homozygous animals. Quantification in **H**. Quantification of MN number and fluorescence intensity in animals expressing *cho-1::mChOpti* in WT and *ceh-20(ay42)* mutants. L4 stage. N > 15. For all quantifications, unpaired two-sided Welch’s t-test was performed, p<.05 = *; p<.01 = **; p<.001 = ***; p<.0001 = ****. Box and whisker plots were used; all data points presented. Box boundaries indicate the 25th and 75th percentile. The limits indicate minima and maxima values. Mean values are highlighted with a black horizontal line.

Animals globally lacking *ceh-20* are larval lethal^57^. To test whether any of the observed Hox effects on terminal identity genes require *ceh-20 (Exd/Pbx1–4)* activity, we employed animals carrying a hypomorphic *ceh-20(ay42)* allele (point mutation in DNA-binding domain)^57^(**Fig. 6A**). In anterior VNC MNs, we observed complete loss of *mig-13::GFP* in *ceh-20(ay42)* animals (L4 stage) (**Fig. 6D**), suggesting that *ceh-20 (Exd/Pbx1–4)* may act as co-factor with *ceh-13 (Lab/Hox1)* and *lin-39* (*Scr/Dfd/Hox3-5*) to co-activate terminal identity gene expression in these cells (**Fig. 6F**). In posterior VNC MNs, we did not observe ectopic *mig-13::GFP* in *ceh-20(ay42)* animals at the L4 stage (**Fig. 6D**). However, examining the same strain at L1, we witnessed ectopic *mig-13* expression in posterior MNs similar to *mab-5(e1239)* mutant animals (**Suppl. Fig. 4**). Although the reason for this stage-specific effect is unclear, these observations suggest that *ceh-20* may be required to activate or repress *mig-13* in a stage-dependent manner (**Fig. 6F**).

In the lumbar MN DA8, we found ectopic *mig-13::GFP* expression in *ceh-20(ay42)* animals at L4 similar to the one observed in *mab-5* mutants (**Fig. 6E**). This was also observed in L1 stage animals (**Suppl. Fig. 4**). Consistently, we found ectopic expression of *itr-1::GFP*, a DA9-specific terminal identity marker (**Fig. 6C**), in DA8 of *ceh-20(ay42)* animals **(Fig. 6E)**, phenocopying the effect seen with the same marker in *mab-5(e1239)* mutants ^19^. Altogether, these data suggest that *mab-5 (Antp/Hox6-8*) and *ceh-20 (Exd/Pbx1–4)* are both required to repress terminal identity gene expression in posterior MNs of the VNC and the lumbar MN DA8.

Besides DA8, *ceh-20* is expressed in three additional lumbar MNs that are cholinergic (DA9, VA12, AS11). However, we observed no effect on *mig-13::GFP* or *itr-1::GFP* expression in these lumbar MNs in *ceh-20(ay42)* animals neither at L1 nor at L4 stages **(Fig. 6E, Suppl. Fig. 4)**. Interestingly, the posterior Hox gene *egl-5 (Abd-A/Abd-B/Hox9-13)* together with the terminal selector *unc-3* are known activators of *mig-13* and *itr-1* in DA9 and VA12 neurons^19^, suggesting that, in these cells, *egl-5* controls MN terminal identity in a *ceh-20* independent manner **(Fig. 6F)**.

Finally, we also examined the effect of *ceh-20* on the expression of the shared terminal identity gene *cho-1/ChT* that is normally expressed in all cholinergic MNs **(Fig. 6C)**. Compared to control animals, *ceh-20(ay42)* mutants showed decreased expression levels of *cho-1* in MNs across the VNC (**Fig. 6G-H)**. However, *cho-1* expression was unaffected in lumbar MNs of *ceh-20(ay42)* mutants, again suggesting that most lumbar MNs do not require *ceh-20* activity for their terminal identity program.

Altogether, these data uncover both PBX-dependent and independent functions for Hox genes in *C. elegans* MNs. In anterior VNC MNs, two Hox genes (*ceh-13, lin-39*), a terminal selector (*unc-3*) and *ceh-20 (Exd/Pbx1–4)* are required for activation of MN terminal identity genes **(Fig. 6F**). In posterior VNC MNs and the lumbar DA8 MN, *mab-5 (Antp/Hox6-8*) and *ceh-20 (Exd/Pbx1–4)* are both required to repress terminal identity gene expression. On the other hand, in other lumbar MNs (DA8, VA12), the posterior Hox gene *egl-5 (Abd-A/Abd-B/Hox9-13)* and the terminal selector *unc-3* activate terminal identity gene expression in a *ceh-20 (Exd/Pbx1–4)*-independent manner (**Fig. 6F**).

### MAB-5 and CEH-20 are continuously required to control MN terminal identity

Prior work demonstrated a role for LIN-39 in the maintenance of MN identity in adult animals^35^, raising the possibility that other Hox factors and PBX proteins may also be continuously required during post-embryonic stages. To test this hypothesis, we specifically focused on MAB-5 and CEH-20, and used the auxin-inducible degron ^58^ system^59^.

First, we used CRISPR/Cas9 to insert an *mNeonGreen::3xFLAG::AID* cassette before the STOP codon of *mab-5* (**Fig. 6A**). Then, we built a strain containing the *mab-5::mNeonGreen::3xFLAG::AID* allele, the *mig-13::GFP* reporter, and a ubiquitously expressing TIR1 transgene PMID. We placed these animals on plates containing auxin starting at L2, a stage by which all MNs have been generated, until the L4 stage (**Fig. 7A**). Compared to animals placed on control (EtOH) plates, we observed efficient depletion of MAB-5::mNeonGreen protein, as well as ectopic *mig-13::GFP* expression in posterior VNC MNs (**Fig. 7B-D**). Although statistically significant, the degree of ectopic expression was smaller compared to *mab-5(e1239)* global mutants, likely due to partial depletion of MAB-5 with the AID system (**Fig. 7D**). From our global mutant and AID analyses, we conclude that *mab-5* is required at late larval stages to repress *mig-13* in posterior MNs.

**Figure 7:**
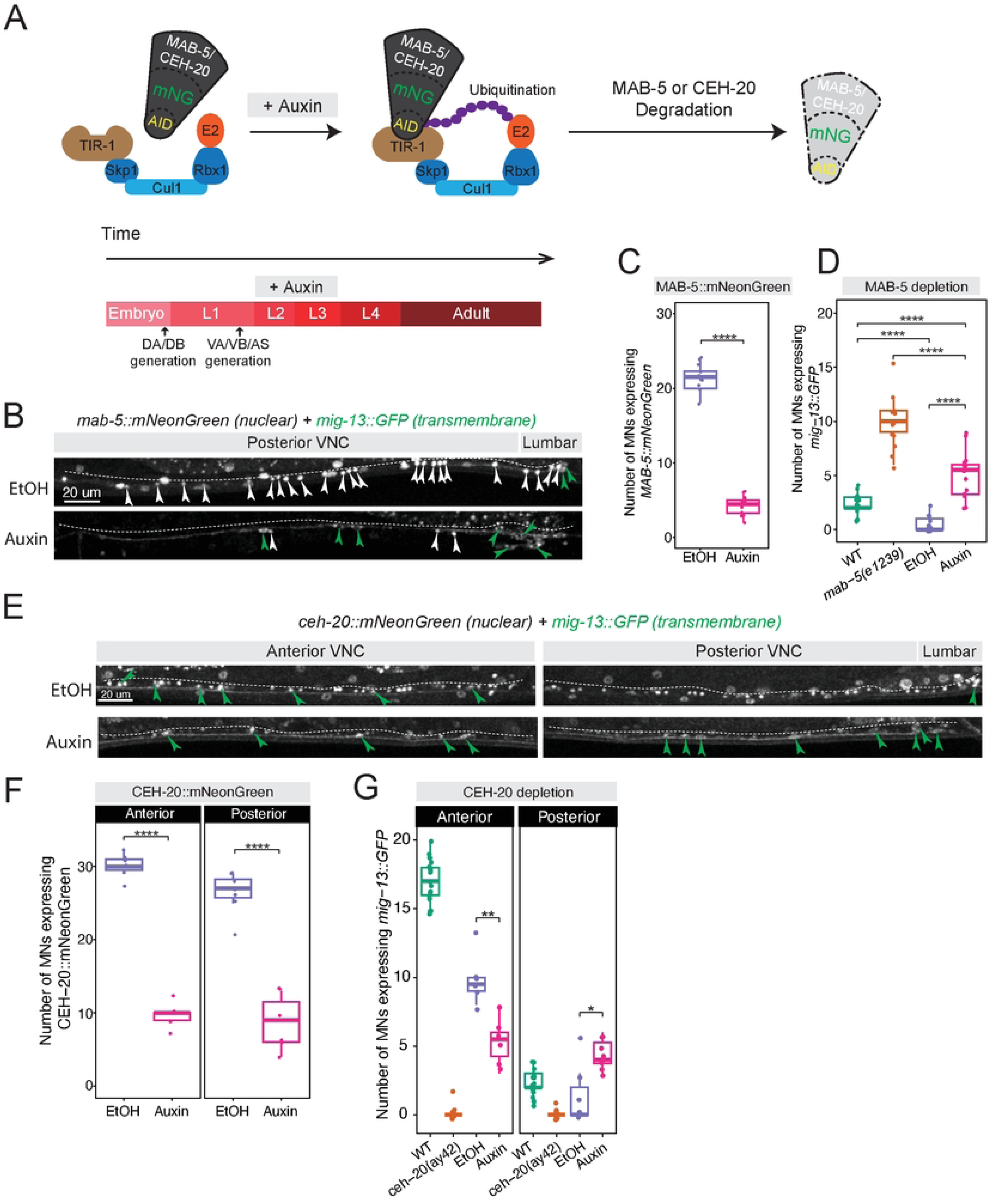
MAB-5 and CEH-20 are required to maintain *mig-13* expression. **A:** For both MAB-5 and CEH-20 depletion, auxin treatment was initiated in animals at the L2 stage, and analysis was performed in L4 animals. **B:** Post-developmental depletion of MAB-5 resulted in ectopic *mig-13::*GFP expression in posterior VNC MNs, like in *mab-5(e1239)* null animals, indicating that MAB-5 is continuously required to repress *mig-13* expression in posterior MNs. **C:** Quantification showing MAB-5 depletion in auxin-treated animals. N= 14. **D:** AID-mediated MAB-5 depletion. Quantification of *mig-13::GFP* in auxin-treated, EtOH (control), and *mab-5(e1239)* animals. **E:** Post-developmental depletion of CEH-20 also resulted in ectopic *mig-13::GFP* expression in posterior VNC MNs, unlike what was observed in *ceh-20(ay42)* L4 animals. **F:** Quantification of CEH-20 depletion in auxin-treated animals. N = 13. **G:** AID-mediated CEH-20 depletion. Quantification of *mig-13::GFP* expression in auxin-treated and *ceh-20(ay42)* animals. For all quantifications, unpaired two-sided Welch’s t-test was performed, p<.05 = *; p<.01 = **; p<.001 = ***; p<.0001 = ****. Box and whisker plots were used; all data points presented. Box boundaries indicate the 25th and 75th percentile. The limits indicate minima and maxima values. Mean values are highlighted with a black horizontal line.

We performed a similar experiment for CEH-20 using our *ceh-20::mNG::3xFLAG::AID* allele. Upon exposure to auxin from the L2 to L4 stage, we observed efficient depletion of CEH-20::mNeonGreen both in anterior and posterior VNC MNs (**Fig. 7F**). Like in *ceh-20(-)* global mutants, auxin-mediated depletion of CEH-20 led to reduced expression of *mig-13::GFP* in anterior MNs (**Fig. 7E-G**), suggesting that *ceh-20* is required at late larval stages to activate *mig-13* in these cells. Conversely, CEH-20 depletion led to increased expression of *mig-13::GFP* in posterior MNs (**Fig. 7E-G**), indicating that *ceh-20* is required late to repress *mig-13* in these cells. Altogether, these findings uncovered MAB-5 and CEH-20 post-embryonic (late larval) requirements for the maintenance of MN terminal identity programs.

### CEH-20/PBX binds terminal identity genes in post-mitotic MNs

To globally assess CEH-20 occupancy at MN terminal identity genes, we interrogated available whole-animal CEH-20 ChIP-seq data^50,60^. These ChIP experiments were performed at the L4 stage, thus including all post-mitotic MNs. Analysis of this dataset revealed strong enrichment of CEH-20 in the *C. elegans* genome, identifying 3,428 unique binding peaks (q-value cutoff of 0.05) (see Methods). Most of the CEH-20 peaks (85.27%) are located between 0 and 2 kb upstream of transcription start sites (TSSs) (**Fig. 8A-B**). The remaining binding events occur between 2 and 3 kb from TSSs (6.86%), at distal intergenic regions (5.28%), and at introns (1.90%). Thus, CEH-20 appears to act primarily at proximal regions (0–2 kb from TSSs) to regulate gene expression.

**Figure 8:**
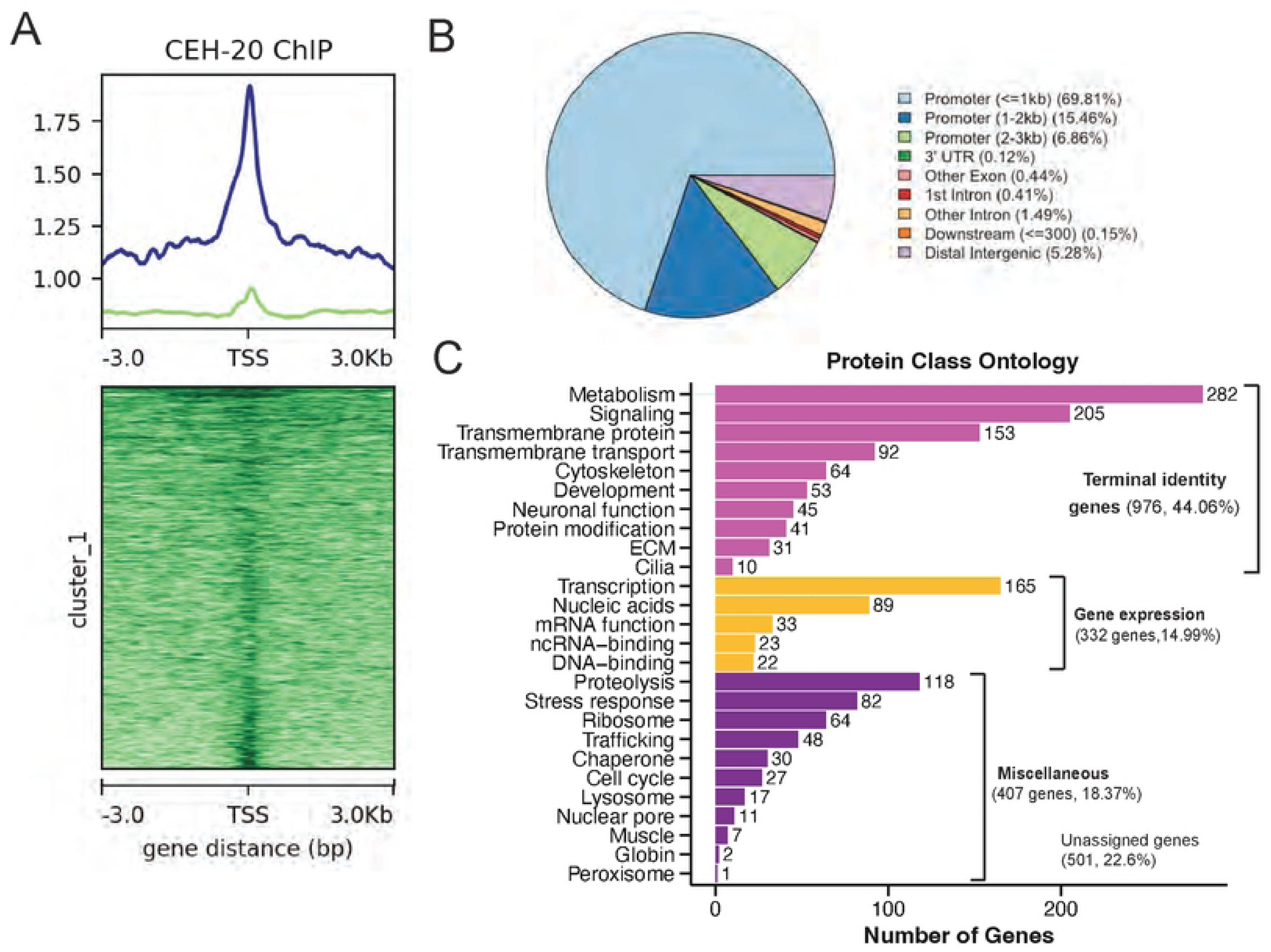
CEH-20/PBX binds terminal identity genes in post-mitotic MNs. **A:** Profile and heatmap of CEH-20 binding relative to TSS. Samples were split into two clusters using K-means^84^. **B:** Pie chart showing genomic distribution of CEH-20 ChIP-Seq peaks. **C:** Gene ontology analysis (WormCat 2.0) of CEH-20 ChIP-Seq target genes that are also expressed in adult VNC MNs.

Next, we annotated the 3,428 CEH-20 ChIP binding peaks and compared the resulting list of 2,968 genes to our scRNA-seq dataset of adult (day 1) *C. elegans* MNs^8^. Out of these 2,968 genes, we identified 2,215 genes (74.63%) bound by CEH-20 to be also expressed in adult *C. elegans* MNs (**Suppl. File 1**), suggesting a prominent role for CEH-20 in MN terminal identity. Next, we conducted Gene Ontology analysis with WormCat 2.0^61^ and found 976 of the 2,215 genes (44.06%) are terminal identity genes (**Fig. 8C**). The second most represented category (287 genes, 14.99%) is genes involved in gene expression (e.g., transcription, mRNA function) (**Fig. 8C**). Taken together, this analysis reveals putative direct CEH-20 target genes in adult *C. elegans* MNs, most of which include terminal identity genes.

### CEH-20/PBX positively regulates Hox factors *ceh-13, lin-39,* and *mab-5*

Our analysis of the CEH-20 ChIP-seq dataset revealed binding peaks at the loci of Hox genes *ceh-13*, *lin-39*, and *mab-5* (**Fig. 9A**), suggesting that CEH-20 may contribute to MN differentiation through Hox gene regulation. To test this hypothesis, we crossed animals carrying transcriptional reporters for *ceh-13 (ceh-13::GFP)*, *lin-39 (lin-39::tagRFP)*, or *mab-5 (mab-5::GFP)* into animals carrying a strong loss-of-function (putative null) allele for *ceh-20 (ok541)* that results in early larval lethality PMID. In each case, we found a significant reduction in the numbers of MNs expressing each Hox gene reporter at L1 (**Fig. 9B-D)**, demonstrating that *ceh-20* activity is required to induce Hox gene expression (*ceh-13, lin-39, mab-5*) in VNC MNs.

**Figure 9:**
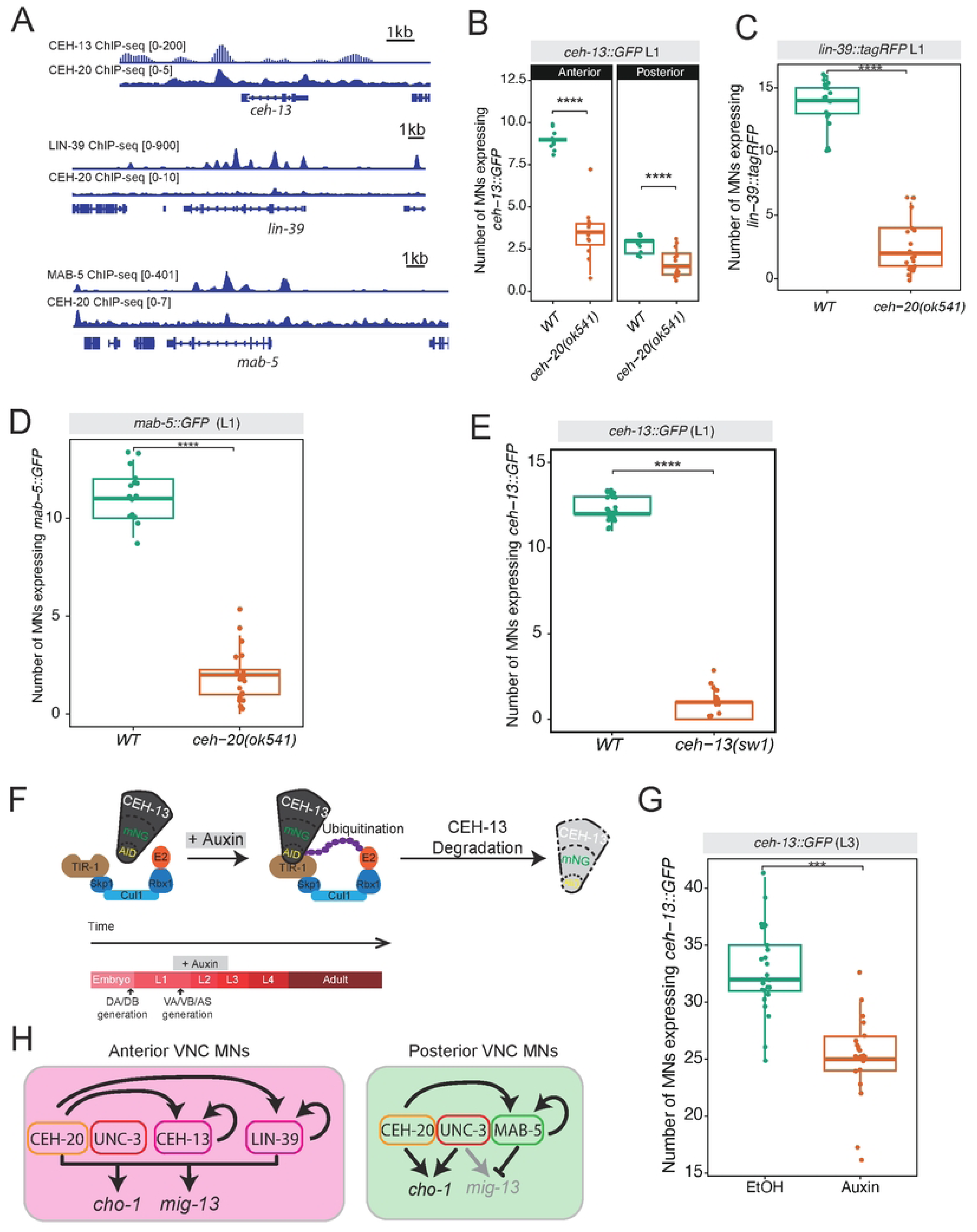
CEH-20 positively regulates three Hox genes in MNs. **A**: Analysis of ChIP-seq shows CEH-20 binding peaks on *ceh-13*, *lin-39*, and *mab-5* loci. Each Hox protein also binds to its cognate locus. **B-D**: The expression of *ceh-13*, *lin-39*, and *mab-5* reporters is reduced in *ceh-20(ok541)* mutants. L1 stage. N = 15. **E**: Reduced expression of *ceh-13::GFP* in *ceh-13(sw1)* mutant animals, suggesting that CEH-13 autoregulates its expression in MNs. L1 stage. N = 15. **F:** The AID system. The E3 ligase complex is composed of Skp1, Cul1, Rbx1, E2. Auxin allows TIR1 to bind the auxin inducible degron, leading to proteasomal degradation of CEH-13 in animals homozygous for the *ceh-13::mNG::3xFLAG::AID* allele (*syb2307*). Auxin treatment was initiated in animals at the L1-2 stage, and analysis was performed in L3-4 animals. **G:** Auxin-induced depletion of CEH-13 protein led to decreased *ceh-13*::GFP (transcriptional reporter) expression in MNs. L3 stage. N=15. For all quantifications, unpaired two-sided Welch’s t-test was performed, p<.05 = *; p<.01 = **; p<.001 = ***; p<.0001 = ****. Box and whisker plots were used; all data points presented. Box boundaries indicate the 25th and 75th percentile. The limits indicate minima and maxima values. Mean values are highlighted with a black horizontal line.

### *ceh-13* (*Lab/Hox1)* maintains its expression in MNs via positive autoregulation

Analysis of available ChIP-seq datasets further showed that CEH-13, LIN-39, and MAB-5 do bind at their own loci (**Fig. 9A**), supporting a model where each of these Hox genes maintains its expression in MNs via positive transcriptional autoregulation. Indeed, this has been previously shown for *lin-39* and *mab-5*^35^. To test whether the model of positive autoregulation also applies to the anterior Hox gene *ceh-13*, we crossed animals carrying a transcriptional *ceh-13::GFP* reporter into *ceh-13 (sw1)* loss-of-function mutants. We found a significant reduction in the number of *ceh-13::GFP*-expressing MNs at L1 (**Fig. 9E**), demonstrating that *ceh-13* gene activity is necessary for its own expression in MNs.

Because the *ceh-13(sw1)* allele removes gene activity starting in early embryo, the above findings do not address whether CEH-13 is required at later life stages to maintain its own expression. To test this, we employed again the AID system using an available *ceh-13(syb2307[ceh-13::3xFLAG::mNG::AID])* allele ^8^ and a panneuronal TIR line (*otTi28*). Continuous exposure to auxin from L1 to L3 stage led to a significant reduction in the number of MNs expressing the transgenic *ceh-13::GFP* reporter (**Fig. 9F-G**), indicating that *ceh-13* is required at larval stages to maintain its own expression. Altogether, these findings strongly suggest transcriptional autoregulation as a positive feedback mechanism for *ceh-13* (*Lab/Hox1)* maintenance in MNs.

## DISCUSSION

Understanding how diverse neuronal subtypes are generated and maintained is a central question in developmental neuroscience. This study provides valuable insights into the transcriptional logic underlying MN subtype diversification along the anterior-posterior (A-P) axis of the *C. elegans* ventral nerve cord. By combining high-resolution expression analysis, functional genetics, and conditional protein depletion strategies, we uncover previously unappreciated roles for Hox transcription factors and the Hox cofactor *ceh-20 (Exd/Pbx1–4)* in MN terminal identity. Our study contributes to our understanding of molecular mechanisms that control neuronal terminal identity in five ways.

First, we describe a new role for the anterior Hox gene *ceh-13* (*Lab/Hox1),* whose function in the *C. elegans* nervous system has remained poorly understood due to early larval lethality of *ceh-13* mutant animals ^63^ ^36,64^. In anterior cholinergic MNs of the nerve cord, our findings support a model where *ceh-13* and the terminal selector *unc-3* act directly to activate multiple terminal identity genes. A prior study in *C. elegans* touch receptor neurons showed that Hox genes can regulate the transcription of terminal selector genes^65^. Therefore, *ceh-13* may also control *unc-3* expression in MNs, in addition to terminal identity genes, though we did not investigate this possibility. Moreover, we show that *ceh-13* positively autoregulates to maintain its expression at later larval stages in anterior MNs. Given the reported positive autoregulation for the mid-body Hox genes *lin-39 (Scr/Dfd/Hox4–5)* and *mab-5 (Antp/Hox6–8)* in *C. elegans* MNs^35^, our findings on *ceh-13* show that positive autoregulation is a broadly applied strategy to maintain Hox gene expression in MNs. Last, the demonstration of *ceh-13* controlling the terminal identity of post-mitotic neurons significantly extends the known repertoire of anterior Hox gene functions in the nervous system. To date, its fly (*Lab*) and vertebrate (*Hox1*) orthologs have only been linked to early events in nervous system development, such as neuroblast proliferation^66^ and rhombomere patterning^67,68^.

Second, we demonstrate that individual Hox genes exert region-specific roles in MN subtype specification by either collaborating with or antagonizing the terminal selector UNC-3. In anterior MNs, *ceh-13* (*Lab/Hox1)* and *lin-39 (Scr/Dfd/Hox4–5)* function together with *ceh-20 (Exd/Pbx1–4)* and UNC-3 to promote terminal identity gene expression. In contrast, in posterior and lumbar MNs, *mab-5 (Antp/Hox6–8)* and *ceh-20 (Exd/Pbx1–4)* repress MN identity programs by antagonizing UNC-3. Intriguingly, inducible protein depletion experiments showed that the repressive function of MAB-5 and CEH-20 extend into late larval stages, suggesting a continuous requirement for HOX and PBX proteins to maintain MN terminal identities throughout life. Such sustained activity expands the classical view of Hox genes as early patterning factors, emphasizing their role in preserving post-mitotic neuronal identity.

Third, our findings show that Hox proteins can operate in both PBX-dependent and independent modes to control MN terminal identity. This is reminiscent of other cellular contexts, where Hox factors can function in either a PBX-dependent or independent manner^23–25,55^. In the context of neuronal development, the PBX-dependent and independent functions of Hox proteins remain poorly defined. A previous study in the mouse spinal cord demonstrated that *Pbx1* and *Pbx3* genes regulate motor neuron (MN) organization and differentiation, acting in either a Hox-dependent or Hox-independent manner depending on the MN subtype^30^. Extending these observations, we find that anterior and posterior MNs of the *C. elegans* ventral nerve cord require terminal selector (*unc-3*), HOX and PBX gene activities to execute their terminal identity programs. On the other hand, the terminal identity of certain lumbar MNs (e.g., DA9, VA12) requires both *unc-3* and the posterior Hox gene *egl-5 (Abd-A/Abd-B/Hox9–13)*, but not *ceh-20 (Exd/Pbx1–4)*. Like our observations in lumbar MNs, posterior Hox genes in *Drosophila* (Abd-B) and vertebrates (Hox9-13) appear to function in a PBX-independent manner ^69–71^.

Fourth, our work uncovers cell-type and region-specific deployment of CEH-20/PBX. While CEH-20 acts as a co-activator with Hox proteins CEH-13 and LIN-39 in anterior MNs, it collaborates with MAB-5 to repress identity programs in posterior VNC and lumbar (DA8) MNs. This dual role of PBX—supporting both gene activation and repression in different MN subtypes—is a striking example of how the same transcriptional cofactor can be redeployed to achieve distinct outcomes on gene expression. Although all Hox proteins possess nearly identical homeodomains and therefore recognize the same, short TAAT DNA motif PMID, *in vitro* studies that assessed HOX paralog binding in the presence of PBX co-factors showed that HOX and PBX proteins not only bind DNA cooperatively, but also recognize a longer motif specific to each HOX paralog^72–75^. Consistent with this idea, LIN-39 and CEH-20 in the *C. elegans* mesodermal lineage are known to bind cooperatively on longer DNA motifs (composed of a HOX and a PBX site) to directly activate gene expression ^76^, a mechanism possibly applicable to *C. elegans* MNs as well.

Finally, PBX’s target genes in the nervous system remain poorly defined. Prior work identified genes involved in neuronal migration and connectivity as PBX targets in mouse spinal MNs PMID. Here, we identify multiple terminal identity genes as CEH-20/PBX targets in *C. elegans* MNs. CEH-20 occupancy at these gene loci supports a direct role in their transcriptional regulation. Further, we find that *ceh-20* is required for the expression of three Hox genes (*ceh-13, lin-39, mab-5*) in MNs. Because all three genes positively autoregulate [this study and ^35^], our findings support a model where each Hox gene binds together with CEH-20/PBX to its cognate locus (**Fig. 9H**), thereby ensuring its maintained expression in MNs. Future work aimed at elucidating the genomic targets and interacting partners of HOX and PBX proteins in specific MN subtypes will shed further light on the logic and evolution of neuronal identity specification.

Taken together, our findings support a model where MN subtype diversity arises through the combinatorial action of terminal selectors and region-specific HOX/PBX modules that differentially activate or repress MN identity programs. Because HOX and PBX proteins are expressed both in invertebrate and vertebrate nervous systems, this modular and context-dependent use of HOX and PBX proteins likely represents a conserved strategy for the control of neuronal diversification.

## ACKNOWLEDGEMENTS

We thank the Caenorhabditis Genetics Center (CGC), which is funded by NIH Office of Research Infrastructure Programs (P40 OD010440), for providing strains. We thank members of the Kratsios lab (Honorine Destain, Filipe Marques) for comments on the manuscript and Stavroula Assimacopoulos for generating plasmids. This work was supported by the Lohengrin Foundation (P.K), a Developmental Biology Training Grant (T32HD055164) to M.P, and an NIH grant (R01NS116365) to P.K.

## AUTHOR CONTRIBUTIONS

M.P, Conceptualization, Data curation, Investigation, Visualization, Methodology, Writing—original draft, review and editing; O.B.E, Y.C., Formal analysis, Validation, Investigation; P. K., Conceptualization, Supervision, Investigation, Funding acquisition, Project administration, Writing— original draft, review and editing.

## DATA AVAILABILITY

All relevant data and processed results supporting the findings are within the manuscript and its Supporting Information files.

## MATERIALS AND METHODS

### *C. elegans* strains and growth conditions

Worms were grown at 20 °C on nematode growth media (NGM) plates seeded with bacteria (OP50, Escherichia coli) as food source^48^. All *C. elegans* strains used in this study are listed in **Suppl. File 2**.

### Targeted genome editing

The endogenous *mig-13* reporter allele *mig-13* (*syb6836 [mig-13::SL2::2xNLS::mScarlet])* was generated by SunyBiotech via CRISPR/Cas9 genome editing by inserting the *2xNLS::mScarlet* cassette immediately before the termination codon of *mig-13*. The endogenous AID-tagged alleles of *mab-5 (syb6730[mab-5::3xFLAG::mNeonGreen::AID])* and *ceh-20 (syb6724[ceh-20::3xFLAG::mNeonGreen::AID])* were also generated by SunyBiotech using CRISPR/Cas9 genome editing.

### Temporally controlled protein depletion

AID-tagged proteins (MAB-5, CEH-13, CEH-20) were conditionally degraded upon exposure to auxin in the presence of TIR1. Animals carrying AID alleles of *mab-5 (syb6730[mab-5::3xFLAG::mNeonGreen::AID])* or *ceh-20 (syb6724[ceh-20::3xFLAG::mNeonGreen::AID])* were crossed to *ieSi57 ([eft-3p::TIR1::mRuby::unc-54 3’UTR + Cbr-unc-119(+)] II; unc-119(ed3) III)* animals, which express TIR1 pan-somatically. Animals carrying the *ceh-13(syb2307[ceh-13::3xFLAG::mNG::AID])* allele (PMID: 38421866) were crossed to *otTi28 [unc-11prom8+ehs-1prom7+rgef-1prom2::TIR1::mTurquoise2::unc-54 3’UTR]* animals, which express TIR pan-neuronally. Natural auxin, indole-3-acetic acid (IAA; Catalog number CAS 87-51-4, Alfa Aesar), was dissolved in ethanol (EtOH) to prepare a 400 mM stock solution. NGM plates containing 4 mM IAA (treatment) and EtOH (control) were prepared, seeded with OP50 bacteria, and allowed to dry for 2-3 days^77^. Worms were transferred onto auxin treatment plates or control plates and kept at 20 °C for the indicated time periods. All experimental plates were shielded from light.

### Microscopy

Worms were anesthetized with 100mM of sodium azide (NaN_3_) and mounted on a 4% agarose pad on glass slides. Images were captured using an automated fluorescence microscope (Zeiss, Axio Imager Z2) or a confocal microscope (Zeiss, LSM 900).

#### Automated fluorescence microscope

Several Z-stack images (minimum thickness ~1 μm) were acquired using the Zeiss Axiocam503 mono and ZEN software (Version 2.3.69.1000, Blue edition).

#### Confocal microscope

Several Z-stack images (minimum thickness ~1 μm) were acquired using the Zeiss Axio Observer 7 and ZEN software (Version 3.8, Blue edition). Representative images are shown following maximum intensity projection of 10–20 μm Z-stacks. Image reconstruction was performed using Image J software^78^.

### Neuron identification

Neurons were identified based on a combination of the following criteria: (i) co-localization with fluorescent markers exhibiting known expression patterns, (ii) invariant cell body position and relative position to other neurons in the preanal ganglion, and (iii) use of the NeuroPAL polychromatic strain^47^.

### Fluorescence intensity quantification

To quantify fluorescence intensity (FI) in individual motor neurons, Z-stack images capturing the ventral nerve cord region and the neurons’ cell bodies were acquired with 0.5-1 μm intervals between stacks. At least 20 worms were analyzed per experimental condition or genotype. The same imaging parameters (e.g., exposure time, temperature) were applied across all samples. Image stacks were then processed, and FI was quantified using Image J (FIJI). All neurons expressing the reporter were counted, and the FI in motor neuron cell bodies was quantified in arbitrary units (a.u).

### ChIP-seq data analysis

CEH-20 ChIP alignment files were downloaded from the ENCODE project website. This ChIP experiment used an anti-eGFP antibody to target endogenously tagged CEH-20 (RW12211 (*st12211[ceh-20::TY1::EGFP::3xFLAG] III)).* BAM files contained reads aligned to the ce11 reference genome. Peak calling was performed with MACS2, applying a minimum q-value cutoff of 0.005 for peak detection ^79^. For visualization, sequencing depth was normalized to 1x genome coverage using bamCoverage from deepTools^80^, and peak signals were visualized in Integrated Genome Viewer ^81^. A heatmap of peak coverage around the CEH-20 enrichment center was generated with NGSplot^82^. The average profile of peaks binding to the TSS region was generated with ChIPseeker^83^. Protein-coding regions were extracted using biomaRt.

### Putative CEH-20 target genes in VNC motor neurons

The top 10,000 highly expressed genes in adult (day 1) motor neurons (measured in transcripts per million, TPM) were extracted from available single-cell RNA sequencing data^8^. This dataset was then computationally compared to a dataset of 2,968 putative target genes at L4 based on CEH-20 ChIP-seq. This comparison generated a new data frame containing 2,215 genes from the scRNA-seq dataset that are also bound by CEH-20. Gene ontology analysis (WormCat 2.0)^61^ was then performed on these 2,215 genes to functionally classify genes based on protein class ontology.

### Statistical analysis and reproducibility

#### Quantification of the number of neurons expressing a reporter gene

Graphs show individual values expressed as mean ± standard error mean (SEM), with each dot representing an individual animal. Statistical analyses were performed using an unpaired t-test (two-tailed). Differences with p < 0.05 were considered significant. Asterisks in figures indicate statistical significance as follows: *p < 0.05, **p < 0.01, ***p < 0.001, ****p<0.0001. Fluorescent microscopy images next to each dot plot show a representative animal. The number of animals (n) for each analysis was determined based on standards in the *C. elegans* field. No data were excluded from the analyses. The Investigators were not blinded to allocation during experiments and outcome assessment.

#### Auxin experiments

For protein depletion, box and whisker plots were used to display all data points, with the whiskers extending to the minimum and maximum values, and the horizontal line within the box representing the mean. Each dot corresponds to an individual neuron. This method also displays each individual value as a point superimposed on the graph. Statistical analyses were performed using an unpaired t-test (two-tailed) with Welch’s correction and p-values were annotated as follows: *p < 0.05, **p < 0.01, ***p < 0.001.

